# Skeletal muscle fibro-adipogenic progenitors of dystrophic mice are insensitive to NOTCH-dependent regulation of adipogenesis

**DOI:** 10.1101/223370

**Authors:** Milica Marinkovic, Francesca Sacco, Filomena Spada, Lucia Lisa Petrilli, Claudia Fuoco, Elisa Micarelli, Theodora Pavlidou, Luisa Castagnoli, Matthias Mann, Cesare Gargioli, Gianni Cesareni

## Abstract

Fibro adipogenic progenitors (FAPs) promote satellite cell differentiation in adult skeletal muscle regeneration. However, in pathological conditions, FAPs are responsible for fibrosis and fat infiltrations. Here we show that the NOTCH pathway negatively modulates FAP differentiation both *in vitro* and *in vivo.* However, FAPs isolated from young dystrophin-deficient *mdx* mice are insensitive to this control mechanism. Nonetheless, factors released by hematopoietic cells restore the sensitivity to NOTCH adipogenic inhibition. An unbiased mass spectrometry-based proteomic analysis of FAPs from muscles of wild type and mdx mice, revealed that the synergistic cooperation between NOTCH and inflammatory signals controls FAP differentiation. These results offer a basis for rationalizing the pathological outcomes of fat infiltrations in skeletal muscle and may suggest new therapeutic strategies to mitigate the detrimental effects of fatty depositions in muscles of dystrophic patients.

**Highlights:**

- Single-cell mass cytometry reveals that *wt* and *mdx* FAPs are in different cell states.
- Activation of the NOTCH signaling pathway negatively regulates adipogenesis of *wt* but not *mdx* FAPs.
- Deep proteomics suggests a mechanism explaining the different sensitivity of *mdx-* FAPs to NOTCH.
- TNF-a stimulation restores the anti-adipogenic effect of NOTCH in *mdx* FAPs.

## Introduction

Skeletal muscle regeneration is a highly orchestrated process involving a variety of mononuclear cell populations that are either resident or attracted to the injured tissue by inflammatory signals (Bentzinger et al., 2013). The interactions of these cell populations contribute to tissue homeostasis. The stability and integrity of the skeletal muscle, however, is altered with ageing or in pathological conditions as a result of a decreased regeneration capacity (Chakkalakal et al., 2012).

A stem cell population residing under the myofiber basal lamina, satellite cells (SC), are the main source of myoblasts during regeneration (Wang and Rudnicki, 2012, Yin et al., 2013). Under physiological conditions SC are quiescent, however, after muscle damage, they are rapidly activated, undergo expansion and myogenic commitment to finally repair the damaged myofibers (Bentzinger et al., 2013, Yin et al., 2013, Wang and Rudnicki, 2012). As a consequence of the exhaustion of the SC stem cell pool in muscular dystrophy patients, the regeneration potential declines and excessive fibrosis and fat infiltrations take place (Rahimov and Kunkel, 2013). Intramuscular adipose tissue is one of the hallmarks of muscular dystrophies, and its extent is a good indicator of disease progression, as it correlates with patient age and clinical stage (Gaeta et al., 2012).

A mesenchymal population of fibro adipogenic progenitors (FAPs), which are located in the interstitial area of the skeletal muscle, positively regulates satellite activation and differentiation (Joe et al., 2010). During muscle regeneration caused by an acute insult, FAPs expand and promote myofiber repair by releasing paracrine factors such as IL-6 and IGF-1 that stimulate satellite cell differentiation (Joe et al., 2010, Farup et al., 2015). Toward the end of the repair process excessive FAPs, which are generated during the expansion phase are removed while the remaining FAPs return to the initial quiescent state (Lemos et al., 2015, Pretheeban et al., 2012, Uezumi et al., 2010, Joe et al., 2010). In pathological conditions, instead of returning to the quiescent state, they rather differentiate causing fibrosis and fat infiltrations (Uezumi et al., 2010, Uezumi et al., 2011, Rodeheffer, 2010).

Despite the established importance of FAPs in both regeneration and degeneration, the signals that regulate the choice between these alternative fates are still poorly characterized. When isolated from the muscle and cultivated *ex vivo* FAPs readily differentiate spontaneously into adipocytes or fibroblasts. This implies that *in vivo* FAP differentiation is negatively controlled by signals from the muscle environment.

As shown by Uezumi and collaborators, non cell-autonomous mechanisms mediated by factors synthetized by regenerating fibers play an important role in limiting adipogenesis during regeneration (Uezumi et al., 2010). Moreover, nitric oxide (NO) has been reported to, affect FAP adipogenic differentiation by down-regulation of peroxisome proliferator-activated receptors gamma (PPARg) (Cordani et al., 2014). Along these lines it has also been proposed that, in acutely damaged skeletal muscle, the balance between the levels of TNF-a and TGFb secreted by infiltrating inflammatory macrophages controls FAP function during regeneration (Lemos et al., 2015). In *mdx* mice the persistence of the anti-apoptotic TGFb signal prevents clearance of the amplified FAPs and favors their fibrotic differentiation. IL-4, secreted by eosinophils has also been reported to inhibit FAP differentiation into adipocytes (Heredia et al., 2013). Taken together these observations underscore the crucial role played by signals from regenerating myofibres or from the inflammatory milieu in controlling FAP differentiation decisions (Bentzinger et al., 2013, Carosio et al., 2011). It is not clear, however, if the shift of FAP fate toward adipogenic differentiation, occurring in dystrophies, can be fully explained by an unbalance of these muscle environmental signals. In alternative, or in parallel, one should consider cell autonomous mechanisms whereby the mutant environment induces a metastable FAP state with a reduced sensitivity to anti-adipogenic signals.

Evolutionarily conserved pathways such as Hedgehog, WNT and NOTCH have been implicated in the regulation of adipogenesis in a variety of experimental systems (Rosen and MacDougald, 2006). In addition, in the muscle system, *in vitro* studies suggest that a direct contact between myotubes and FAPs negatively affect FAP adipogenic differentiation (Uezumi et al., 2010, Huang et al., 2014). We focused on the NOTCH pathway since i) it is activated by cell-cell contact (Andersson and Lendahl, 2014) ii) it is a known regulator of stem cells quiescence (Koch et al., 2013) iii) it plays a fundamental role in muscle regeneration (Mourikis and Tajbakhsh, 2014).

Here we report that NOTCH also modulates FAP adipogenesis and that this control mechanism is compromised in FAPs from dystrophin deficient, *mdx,* mice, a model of Duchenne Muscular Dystrophy (DMD). Our results support a model whereby the integration of NOTCH with other anti-adipogenic signals plays an important role in the regulation of FAP adipogenesis in a healthy muscle and in an *mdx* background.

## Results

### 1. FAPs are a heterogeneous cell population and the abundance of phenotypically distinct subpopulation vary in FAPs from regenerating muscles

Fibro adipogenic progenitors were purified from muscle mononuclear cells by the MACS microbead technology as CD3Γ/CD45Vα7-integrin^-^/SCA1^+^ cells (from now on, FAPs). FAPs isolation strategy, characterization and evaluation of differentiation potential are presented in Supplemental Figure 1. We first asked whether FAPs from mdx dystrophic mice can be phenotypically discriminated from wild type. *mdx* FAPs have different differentiation potentials *in vivo* and *ex vivo* when compared to wild type ((Mozzetta et al., 2013)and our unpublished observations). We found that this phenotypic difference is reflected by differences in the surface protein expression profile as revealed by mass cytometry (Fig. 1). For this analysis we isolated and compared the antigen profiles of FAPs from 6 weeks old wild type (*wt*) and *mdx* mice. At this age the hind limb muscles of *mdx* mice are in a robust regeneration phase (Pastoret and Sebille, 1995). We also included in the analysis FAPs from a second model of muscle regeneration obtained by purifying mononuclear cells 3 days after cardiotoxin (*ctx*) injury of mice of comparable age (Fig. 1A). Each condition (wt, *mdx, ctx*) was analyzed in three biological replicates. The nine samples were each tagged with palladium isotopes, using a bar-coding protocol (Bodenmiller et al., 2012). Cells were then combined in a single suspension and tagged with a heavy atom labeled antibody panel, including 12 antibodies, six of which were against surface antigens (Fig. 1 and 2) (Table S1). While most of the antigens showed a quasi modal distribution of the intensities, with minor, albeit significant, shifts in average intensities (see for instance CD140b in Fig. 1B), SCA1, CD34 and CD146 intensity distributions were clearly bimodal (Fig. 1B). In addition to a low expression peak characterizing the wild type FAP preparation, the *mdx* and *ctx* preparations displayed a second peak of cells expressing higher levels of CD34 and/or SCA1 (Fig. 1B), the second population being more numerous in the *ctx* FAP preparations. The expression of SCA1 and CD34 were highly correlated defining two FAP sub-populations with high or low expression of both antigens (circled in green and yellow in Fig. 1C). The populations expressing anticorrelated levels of SCA1 and CD34 were negligible. The SCA1^L^CD34^L^ and SCA1^H^CD34^H^ subpopulations characterize the *wt* and *ctx* preparations respectively while the *mdx* FAP preparation contained both sub-populations with an approximately equal number of cells (Fig. 1D).

**Figure 1.**
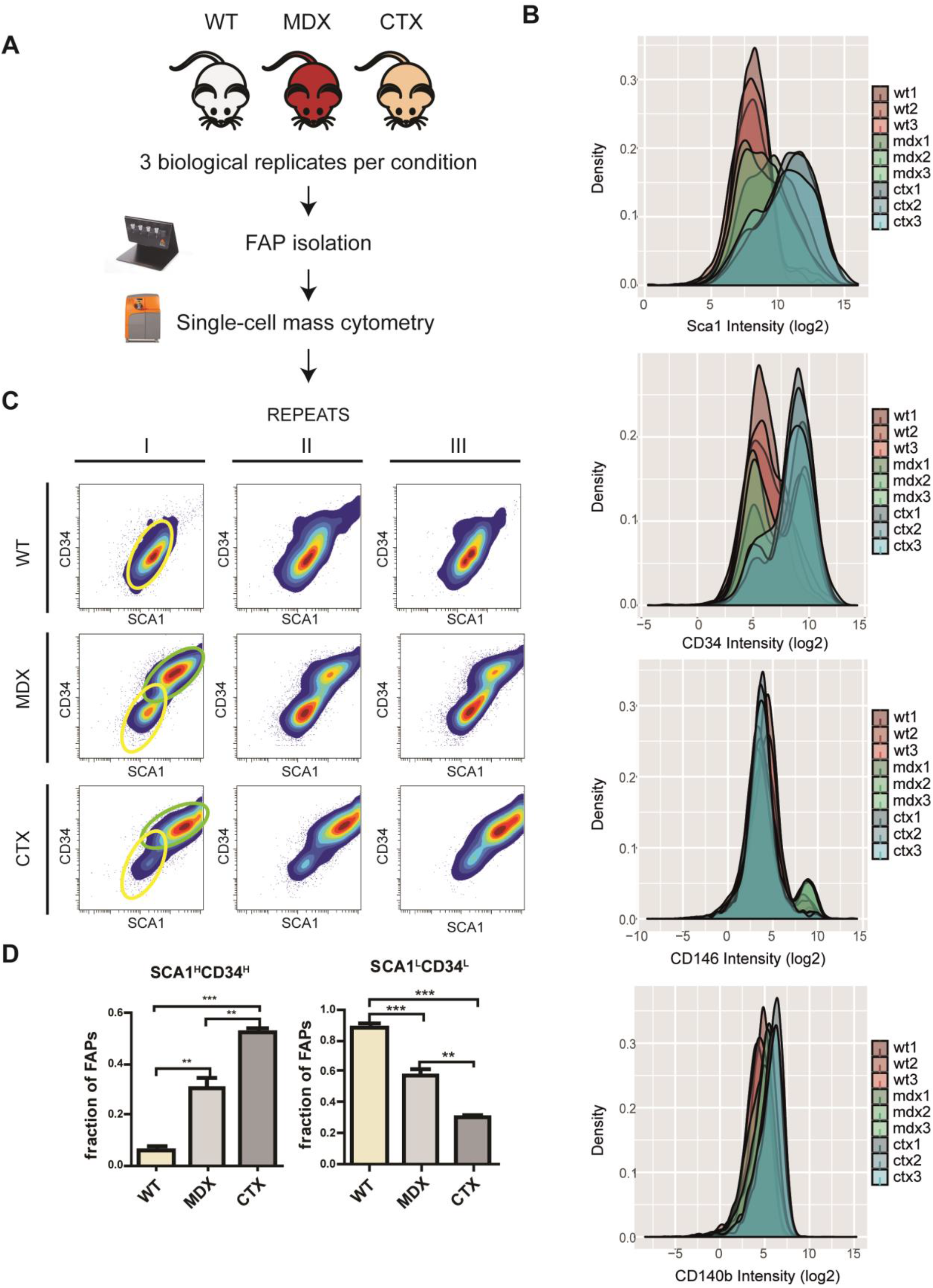
Mass cytometry analysis of *wt, mdx* and *ctx* FAPs. FAPs were purified using MACS microbead technology from hind limb muscles of six-weeks old *wt, mdx* and *ctx* mice. The different cell samples were bar coded by labeling with different proportions of three palladium isotopes. The bar-coded cell samples were pooled and incubated with 12 metal labeled antibodies (SCA1, CD34, CD146, CD140b, CD31, CXCR4, Cas3, Vim, pSta1, TNF-a, pCreb, pStat3) and analyzed by mass cytometry in a Cytof2 instrument. A) Schematic representation of the experimental design. B) Intensity distribution of the signals of four different antigens (SCA1, CD34, CD146 and CD140b). C) Contour maps of the scatter plots showing the correlations between SCA1 and CD34 expression in the three biological repeats for the three conditions. Colors represent cell densities: red=high blue=low. The two different identified sub-populations are outlined by green (SCA1^H^,CD34^H^) or a yellow (SCA1^L^CD34^L^) ovals D) Bar plots representing the abundance of the two subpopulations expressing different levels of SCA1 and CD34. Statistical significance was estimated by the ANOVA test (** p<0.01, *** p<0.001).

The mass cytometry multiparametric data was analyzed using the viSNE algorithm (Amir el et al., 2013). By projecting the single cell multiparametric data onto a two-dimensional plane we could identify FAP subpopulations whose abundance changed in the different conditions. This confirms that the CD3^-^/CD45^-^/α7-integrinVSCA1^+^ cells, form a rather heterogeneous population whose states are influenced by the conditions of the muscle environment (Fig. 2A). For the sake of discussion, we identify 5 different sub-populations expressing characteristics antigen levels (A to E in Fig. 2B). Striking is the difference in the shape of the viSNE plot when *wt* FAPs are compared to *mdx* and *ctx* FAPs. While populations B and C, expressing relatively low levels of SCA1 and CD34, are more abundant in the *wt* preparation and hardly present in the *ctx* sample, subpopulation D and E are consistently more numerous in all the biological replicates of the *mdx* and *ctx* preparations (Fig. 2A). Sub-population D and E both express significantly higher levels of the SCA1 and CD34 antigens but are here considered distinct because population E in addition to expressing even higher levels of SCA1 also expresses higher levels of CXCR4 and TNF-a (Fig. 2B) Subpopulations D and E populate the FAP samples from *mdx* and *ctx* mice, being more numerous in *mdx* and *ctx* respectively. Overall *mdx* and *ctx* viSNE plots are qualitatively similar. However, striking in this respect is the identification in *mdx* muscles of subpopulation A characterized by the expression of the perycitic marker CD146 (Crisan et al., 2012). This subpopulation is hardly present in the *ctx* preparation.

**Figure 2.**
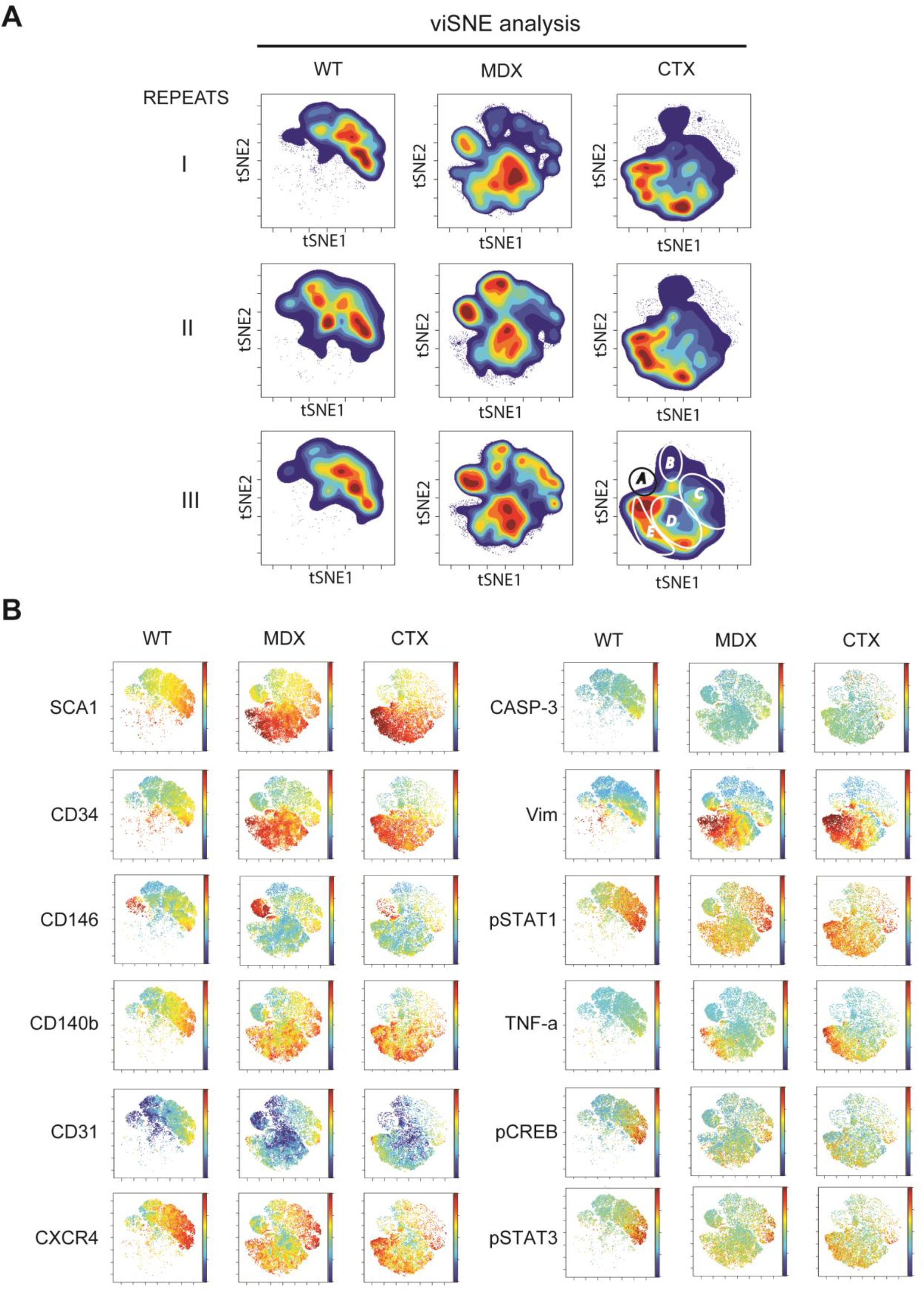
viSNE analysis of the multiparametric data of the FAP preparations. The single cell multiparametric data obtained in the mass cytometry experiment described in Figure 2 was projected onto a two-dimensional space by applying the viSNE algorithm (Amir el et al., 2013). A) Contour representations of the viSNE maps In the nine different cell preparations. The maps are colored according to the cell density in the specific area of the tSNE two dimensional maps (red= high density, blue= low density). Regions of interest identifying cell subpopulations that characterize the different biological conditions are circled (A to E in the top left panel) B) Dot representations of the viSNE maps where each dot represents a cell. In the different maps dots are coloured according to the intensity of expression of the 12 antigen readouts.

In conclusion, when the antigen expression profiles of FAPs from mdx or *ctx* mice are compared with wild type by single cell analysis, they show remarkable and reproducible differences. The clearly recognizable viSNE two dimensional plots suggests that the FAPs are in different cellular states depending on the muscle environment experienced *in vivo.* We speculated that these different states could underlie a different ability to respond to differentiation stimuli.

### 2. Manipulation of NOTCH signaling pathway perturbs FAP differentiation into adipocytes *in vitro*

The single cell analysis of FAPs from three experimental conditions (wt, *ctx* and *mdx* mice) detected a shift in the abundance of populations expressing different levels of FAP antigens. Notwithstanding these different antigen profiles the three FAP preparations efficiently differentiate into adipocytes when isolated from the muscle environment and cultivated *ex vivo* (not shown). We were interested in investigating signals keeping FAPs adipocytic differentiation potential under control *in vivo* and to ask whether the different cell states observed by mass cytometry analysis have the potential to affect, in a cell autonomous manner, their sensitivity to the inhibitory signals provided by other cell types in the muscle environment. In order to identify the regulatory mechanisms that may control FAP differentiation, we investigated six signals that activate or inhibit three key pathways, NOTCH, WNT and Hedghog, that were reported to modulate adipogenesis in mesenchymal progenitor cells (Rosen and MacDougald, 2006). Our analysis revealed that that the modulation of NOTCH pathway had the most striking effect on FAP adipogenesis, without affecting cell proliferation and survival (Fig. S2).

Next, we isolated FAPs from wild type mice and treated them with different concentrations of the γ-secretase inhibitor DAPT (Fig. 3A) during differentiation in growth medium (DMEM+ 20% FBS) changing the medium every 48h for 8 days (Fig. 3B). We observed a remarkable dose-dependent increase of the fraction of cells stained with Oil red O when compared to untreated controls (Fig. 3 C, E). A similar trend was observed when adipogenic commitment was observed by staining with antibodies against PPARg, the adipogenesis master regulator gene (Fig. 3 C, E). The average number of nuclei did not significantly change in treated groups compared to controls (Fig. 3D). These results show that γ-secretase negatively controls adipogenesis. Since γ-secretase is essential for NOTCH activation, this observation is consistent with the hypothesis that NOTCH is involved in the regulation of adipogenesis. We conclude that FAPs, by synthesizing both a NOTCH ligand and a NOTCH receptor are able to limit their own differentiation by an autocrine mechanism.

**Figure 3.**
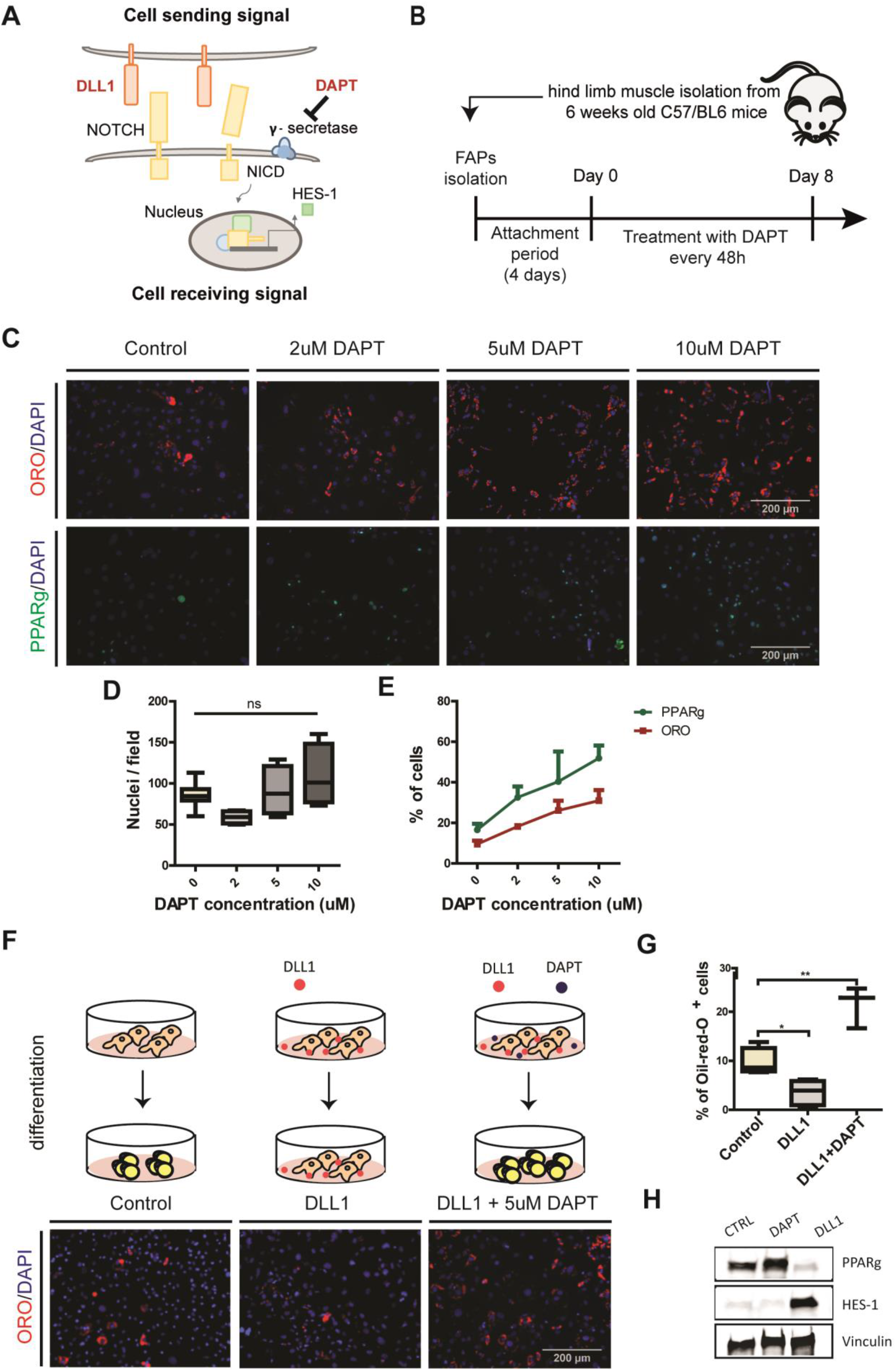
Perturbation of the NOTCH pathway modulates FAP differentiation into adipocytes. A) Simplified model of canonical NOTCH signaling. DAPT (N-[N-(3, 5-difluorophenacetyl)-L-alanyl]-S-phenylglycine t-butylester) is a synthetic inhibitor of γsecretase (B) Schema of the experimental procedure. C) FAPs were isolated from C57BL/6 mice, cultivated in DMEM+20%FBS and treated upon attachment with 2 μM, 5 μM and 10 μM of DAPT every 48h for 8 days. Adipocytes were detected by Oil red O stain or PPARg, and nuclei were counterstained with DAPI. (D) The average number of nuclei are presented as a box plot (n=4). (E) Line graph showing the percentage of Oil red O and PPARg positive cells represented as average ± SEM of four different experiments (n = 4). F) FAPs isolated from hind limb muscles of C57BL/6 mice were plated on DLL1-Fc or IgG2A-Fc (control) coated surface in DMEM+20%FBS. Additionally FAPs were plated on DLL1-Fc coated surface and treated with 5μM DAPT every 48h for 8 days. Adipocytes were detected with Oil red O stain and nuclei counterstained with DAPI. (G) The percentage of Oil red O positive cells of at least three different experiments are shown in the graphs. All quantifications were done using the CellProfiler software. Box plots show median and interquartile range with whiskers extended to minimum and maximum values.Statistical significance was evaluated by the ANOVA test (*p<0.05, **p<0.01). H) Western blot analysis of HES-1 and PPARg protein levels in control, DAPT and DLL1 treated cells. Proteins (30ug) from FAP whole cell lysates were separated by electrophoresis on an SDS acrylamide (4–15 %) gel and blotted onto nitrocellulose paper. Vinculin was used as a loading control (n=3).

To further support the role of NOTCH signaling, we next asked whether activation of NOTCH signaling by exposure to the ligand DLL1 affects FAP adipogenic differentiation. FAPs isolated from wild type mice were plated on DLL1-Fc coated surface and cultivated in growth medium for 8 days. Plates coated with IgG2A-Fc were used as a control. One sample was additionally treated with 5μM DAPT every 48h. FAPs seeded on plates coated with DLL1, have a significant impairment of adipogenic differentiation. However, this inhibition was overridden by exposing cells to 5μM DAPT (Fig. 3 F-H). The activation of NOTCH pathway and the inhibition of adipogenesis was monitored by western blot analysis of HES-1 and PPARg expression, a down-stream NOTCH target and the master transcription factor of adipogenesis respectively (Fig. 3H).

### 3. Myotubes inhibition of FAP adipogenesis is NOTCH dependent.

The crosstalk between different cells in the skeletal muscle stem cell niche is crucial for the regulation of muscle homeostasis (Bentzinger et al., 2013). In order to understand the cellcell interaction controlling FAPs adipogenesis, we aimed to identify the muscle resident cell type with the potential to activate the inhibitory stimulus mediated by NOTCH pathway. Myotubes can inhibit FAP differentiation when co-cultured *ex vivo* (Huang et al., 2014; Uezumi et al., 2010). We confirmed that seeding purified muscle satellite cells (SC) and FAPs in co-culture, in a 1:1 ratio, inhibits lipid droplets formation even in conditions favoring adipogenic differentiation (Fig. 4 B-D). SC and FAPs were isolated from muscles of C57Bl/6 mice and plated in co-culture conditions in satellite cells growth medium. Upon attachment, cells were propagated for two days in the same medium. In order to insure adipogenic differentiation of FAPs, cells were first stimulated with adipogenic induction medium for 48h, which was then replaced with adipogenic maintenance medium for 4 additional days (Fig. 4A). As already reported by Uezumi et al (2010) we confirmed that adipogenic inhibition requires the direct contact between the two cell populations, as co-culturing the two cell types in separate compartments, by using transwell inserts with 1μm porous membrane adipogenesis is not affected (Fig. 4 B-D). Adipocyte differentiation in co-cultures was estimated by monitoring the fraction of Oil red O positive cells (Fig. 4C) and by measuring the levels of adiponectin secreted in the medium (Fig. 4D).

**Figure 4.**
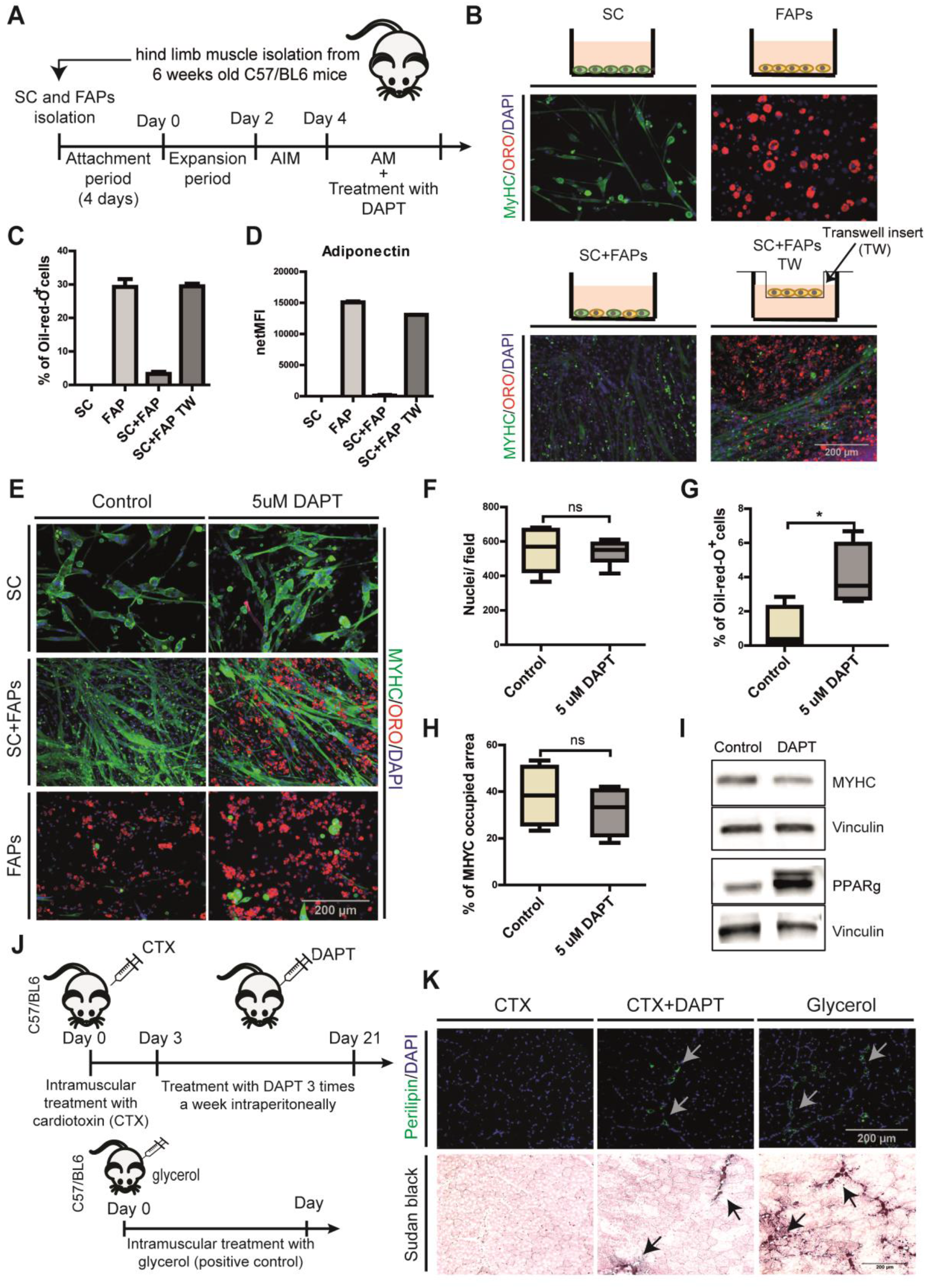
Myotube inhibition of FAP adipogenesis is dependent on NOTCH activity. A) Schema of the experimental procedure. B) Satellite cells and FAPs were cultured separately or cocultured (directly or in transwell) in satellite cells growth medium for 2 days, followed by 2 days in adipogenic induction medium (AIM) and 4 days in adipogenic maintanance medium (AM). Cells were stained with anti MYHC antibodies, Oil red O and DAPI. C) The percentage of cells stained with oil red O and D) the amount of adiponectin secreted in the culture medium was quantitated and reported as bar graphs showing average±SEM (n=2). E) Satellite and FAPs were cocultured and treated with 5μM DAPT every 48h hours. (F,G,H) Box plots representing the average number of nuclei per field, the fraction of cells stained with Oil red O and the fraction of pixels stained with anti MYHC antibodies respectively. Quantifications were done using the CellProfiler software. Box plots show median and interquartile range with whiskers extended to minimum and maximum values (n=4). Statistical significance was evaluated by the Student’s t-test (*p<0.05). I) Proteins (30ug) from the coculture cell lysates were separated by SDS PAGE. Western blot analysis of MYHC and PPARg expression in control and treated co-cultures. Vinculin was used as a loading control, (n=3). J) Schema of *in vivo* experiment. Six weeks old wild type, C57BL/6, mice were injected with cardiotoxin (10μM) and starting the 3^rd^ day after injury were treated with 30 mg/kg DAPT or vehicle, intraperitoneally 3 times a week until day 21. Mice treated intramuscularly with glycerol 50% (Hank’s Balanced Salt Solution, v/v) were used as a positive control for intramuscular adipocyte accumulation. K) Representative TA sections of control and mice treated with DAPT or glycerol stained with perilipin and DAPI or sudan black (n control=2 mice, n DAPT= 2 mice).

Given that in skeletal muscle the delta-like-ligand 1 (DLL1) is expressed on the cell membrane of satellite cells and myofibers (Bi et al., 2016, Conboy et al., 2003, Conboy and Rando, 2002, Kuang et al., 2007), and since NOTCH signaling is transmitted via cell-cell contact, we investigated if the inhibition observed in co-culture was mediated by activation of NOTCH signaling. To this end we treated the SC-FAPs cocultures with 5μM DAPT every 48h (Fig. 4A). When FAPs were treated with DAPT in co-culture conditions adipocytes differentiation was observed, while satellite differentiation and myotube formation was unaffected (Fig. 4 E-I). In addition, we did not observe a significant difference in the number of nuclei in controls and DAPT treated co-cultures, suggesting that proliferation was also not affected (Fig. 4F). These results were confirmed with western blot analysis for myosin heavy chain (MYHC) and PPARg expression in control and DAPT treated co-cultures (Fig. 4 I). Taken together these results suggest that satellite cells, or satellite cell-derived myotubes, inhibit FAP differentiation by activating NOTCH.

We next asked whether NOTCH has any role in limiting adipogenesis *in vivo* during muscle regeneration. Cardiotoxin-injured mice were treated by intraperitoneal injection of DAPT (30 mg/kg, three times per week) starting 3 days after the muscle injury for 3 weeks. A mild fatty tissue infiltration was revealed by perilipin immunostaining and by Sudan Black stain only in the group treated with DAPT (Fig. 4 J, K). As a positive control, we used mice treated with glycerol, a common model for skeletal muscle fat infiltration (Pisani et al., 2010).

Overall our observations support a novel role of NOTCH in muscle regeneration, which is not only limited to modulation of satellite cell activation and differentiation, but also participates in the control of FAP differentiation. The mild pro-adipogenic effect observed after NOTCH inhibition is consistent with the existence of multiple control systems that cooperate to limit adipogenesis.

### 4. FAPs isolated from *mdx* mice are insensitive to adipogenic inhibition by NOTCH activation

Building on the observation that NOTCH suppresses FAPs adipogenesis in wild type mice, we asked if the same mechanism is responsible for restraining FAPs from differentiating into adipocytes in young *mdx* mice. FAPs were isolated from 6 weeks old, *mdx* mice and seeded on plates coated with the NOTCH ligand DLL1. Strikingly, *mdx* FAPs did not show any significant variation in proliferation or differentiation when compared to control cells (Fig. 5 A-C). This result suggests that, differently from wild type, FAPs from *mdx* mice are insensitive to NOTCH inhibition.

**Figure 5.**
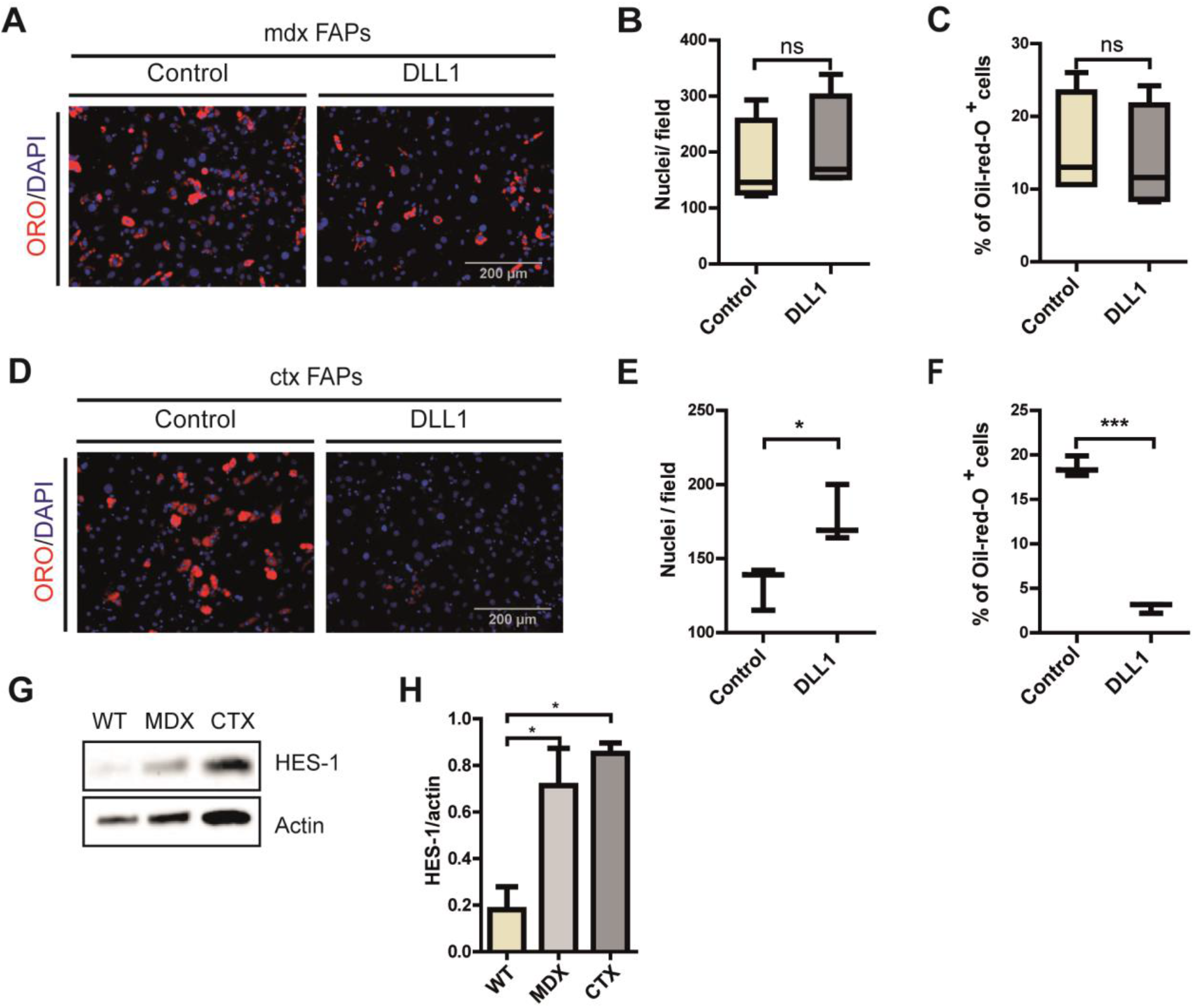
mdx FAPs are unresponsive to DLL1 activation of NOTCH signallng. A) FAPs isolated from young (6 weeks old) *mdx* mice were plated on DLL1-Fc or IgG2A-Fc (control) coated surface in growth medium for 8 days. Adipocytes were stained with Oil red O and nuclei with DAPI. (B, C) The average number of nuclei and percentage of adipocytes are presented as box plots (n=4). D) FAPs isolated from *ctx*-treated C57BL/6 mice 3 days after treatment, were plated on DLL1-Fc or IgG2A-Fc (control) coated surface in growth medium for 8 days. Adipocytes were stained with Oil- ed O and nuclei with DAPI. E, F) The average number of nuclei and percentage of adipocytes are presented as a box plot (n=3). All quantifications were done using the CellProfiler software.Box plots show median and interquartile range with whiskers extended to minimum and maximum values. Statistical significance was evaluated by the Student’s t-test (*p<0.05). G) Expression of HES-1 in FAPs from three conditions (*wt, mdx, ctx*) was evaluated. Proteins were isolated from freshly isolated FAPs from *wt, mdx* and *ctx*-treated mice and the expression of HES-1 was detected by immunoblotting and H) quantified by densitometric analysis (n=3). Actin is used as a loading control. Statistical significance was evaluated by the ANOVA test (*p<0.05, *** p<0.001).

Given that young *mdx* mice are characterized by repeated regeneration-degeneration cycles (Grounds et al., 2008), we entertained the hypothesis that this decreased sensitivity to NOTCH ligand could be a consequence of cell state perturbation due to chronic inflammation exposure. To address this hypothesis, 6 weeks old *wt* mice were treated with cardiotoxin (*ctx*) to induce inflammation and regeneration in a wild type muscle. Three days post-injury FAPs were isolated, as at this point they reach their proliferation peak (Lemos et al., 2015). However, when seeded on DLL1 coated surface FAPs prepared from these regenerating muscles failed to differentiate and were as sensitive to NOTCH inhibition as FAPs prepared from untreated muscles (Fig. 5 D-F).

These results, highlights a differential sensitivity of wild type and *mdx* FAPs to NOTCH mediated modulation of differentiation and may provide a molecular explanation of the observed fat infiltrates in aging dystrophic organisms.

Interestingly, the HES-1 protein, the product od a down-stream target of the NOTCH pathway, is more expressed in mdx rather than wild type FAPs, suggesting that the canonical NOTCH pathway is functional in *mdx* FAPs (Fig. 5 G, H). Upregulation of Hes-1 is observed also in *ctx* FAPs suggesting that the NOTCH signaling pathway is activated in FAPs upon injury. We surmise that the insensitivity of *mdx* FAPs to the anti-adipogenic effect of NOTCH activation is not a consequence of a defect in some components of the canonical NOTCH signaling machinery but rather that the inhibition of adipogenesis is the result of a cross-talk between the NOTCH pathway and an as yet unidentified molecular mechanism which is altered in dystrophic FAPs.

### 5. Remodeling of the MDX FAPs proteome

To elucidate the molecular mechanisms underlying the different sensitivity of *mdx* FAPs to NOTCH modulation, we performed deep mass spectrometry (MS)-proteomic profiling of FAPs freshly isolated from hind limb muscles of wt, *mdx* and *ctx* mice. We applied the recently developed iST proteomics workflow (Kulak et al., 2014), combined with label-free LC–MS/MS analysis (Fig. 6A), and processed the results in the MaxQuant environment (Cox and Mann, 2008, Cox et al., 2011). All experiments were performed in at least biological triplicates, revealing high quantitation accuracy and reproducibility, with Pearson correlation coefficients between 0.85 and 0.95 (Fig. S3A). This approach enabled the quantification of about 7,000 proteins (Table S2) with abundance spanning a range covering 7 orders of magnitude. As previously described in cell lines, structural proteins and proteins in basic cellular machineries are much more abundant than regulatory proteins, such as transcription factors (Fig.S3B). To investigate whether our large-scale proteomic data could enable unsupervised classification of *mdx, ctx* and *wt* FAPs, we performed principal component analysis (PCA) of approximately 4000 proteins quantified in at least 50% of our experimental conditions. Remarkably, PCA segregated the three biological conditions into three main clusters (Fig. 6B) confirming little variability in the proteomic profile of biological replicates. To investigate the possible functional consequence of proteome remodelling in *mdx* and *ctx* FAPs, we focused on the 2041 proteins significantly modulated across the three different experimental conditions (T-Test, FDR<0.05) (Fig. 6C). This approach enabled the identification of three groups of significantly modulated proteins: i) *mdx*-specific proteins; ii) *ctx*-specific proteins; iii) proteins similarly modulated in *mdx* and *ctx* FAPs when compared to *wt* FAPs. Next we investigated whether these groups of proteins were enriched for specific biological processes or pathways. Interestingly we found that proteins involved in cell cycle and DNA replication were upregulated in both *mdx* and *ctx* FAPs. Consistent with this observation, FAPs are known to be activated and proliferate after chronic and acute muscle damage (Joe et al., 2010). Additionally, we observed that proteins involved in TCA cycle and fatty acids metabolism were down-regulated in *mdx* and *ctx* FAPs. This suggests that, similarly to satellite cells, FAPs activation during the muscle regeneration process may experience a metabolic switch (Ryall et al., 2015).

**Figure 6.**
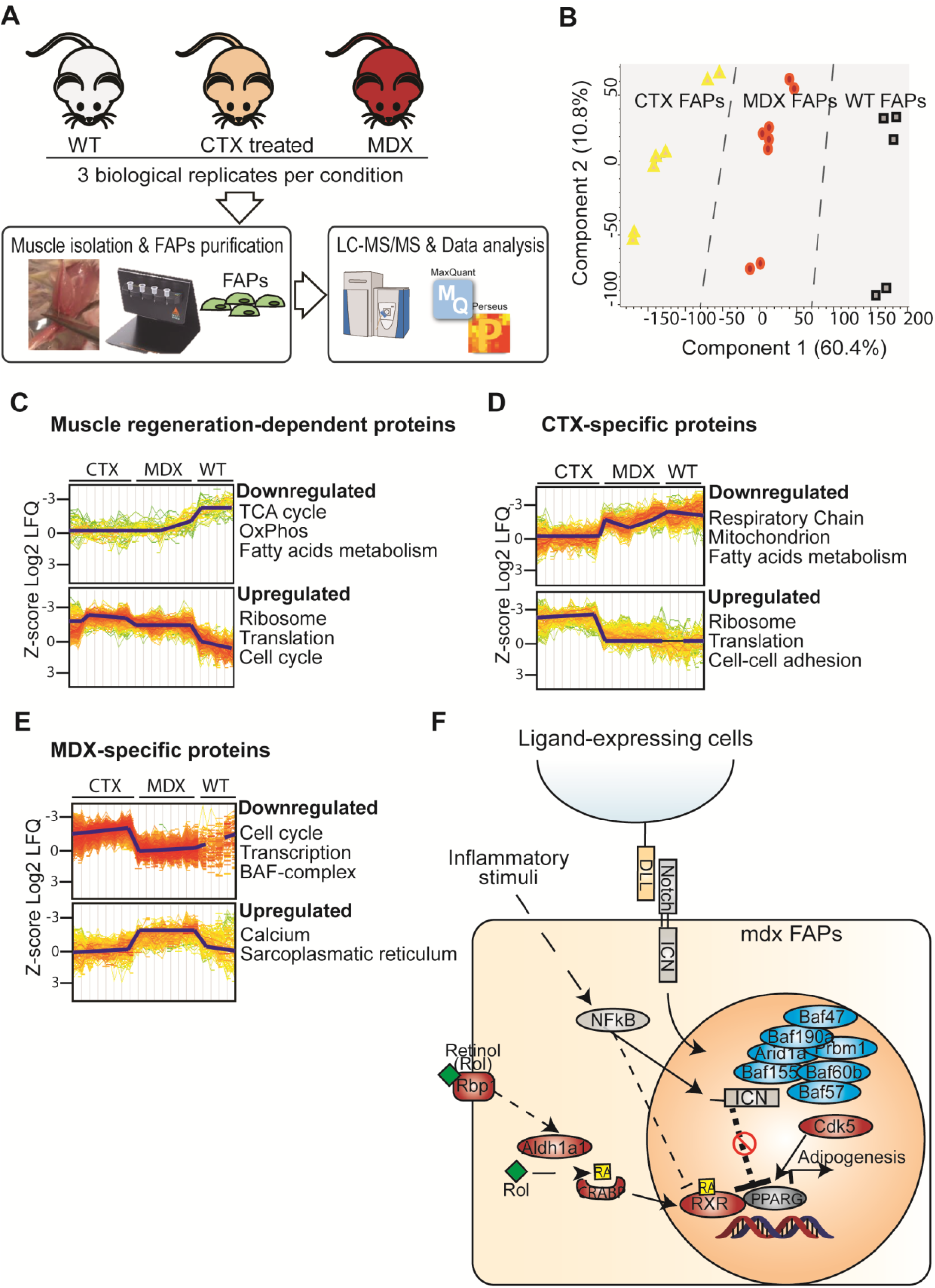
Deep-MS based proteomic approach reveals mechanisms underlying the different sensitivity of mdx FAPs to NOTCH stimulation. A) Schematic representation of the experimental strategy applied to analyze the proteome of FAPs isolated from wt, mdx and cardiotoxin-treated mice. B) Principal Component Analysis (PCA) indicate good separation between wt, *mdx* and *ctx* FAPs. C-D-E) GO BPs and pathways enrichment analysis in selected clusters of significantly modulated proteins (T-test, FDR<0.07). F) Model recapitulating the mechanisms identified by the MS-based proteomic approach through which mdx FAPs may escape the NOTCH-dependent inhibition of adipogenesis. The proteins that are up- or down-regulated in mdx FAPs when compared to *ctx* FAPs are in red and blue respectively

Next to identify mechanisms that would explain the reduced sensitivity of *mdx* FAPs to the anti-adipogenic signals triggered by NOTCH, we investigated whether the 600 proteins that are specifically modulated in FAPs from mdx mice were functionally connected to the NOTCH pathway and to the adipogenesis regulatory network. The overlay of our proteomic data onto the NOTCH and adipogenesis pathways, compiled from the literature information curated in the SIGNOR database (Perfetto et al., 2016) highlighted different mechanisms that may underlie the diverse response of *mdx* FAPs to NOTCH-dependent adipogenic signals: i) the down-regulation of many components of the SWI/SNF chromatin remodeling complex, known to positively modulate the transcriptional activity of NOTCH (Yatim et al., 2012); ii) the upregulation of the CDK5, kinase, which phosphorylates and promotes the transcriptional activity of the adipogenesis master gene, PPARg (Choi et al., 2010); iii) the up-regulation of proteins involved in retinol metabolism including the retinoic receptor RXRA, which forms a transcriptionally active heterodimer with PPARg, driving an adipogenic differentiation program (de Vera et al., 2017)

Given the importance of retinoic acids in adipogenic control (Berry et al., 2012), we focused out analysis on the *mdx*-dependent upregulation of the RXRA receptor. Interestingly both RXRA and NOTCH pathways are regulated by the activation of NFKB, a master regulator of the inflammatory response. Specifically, while NFKB inhibits the RXRA-PPARg complex through the upregulation of the HDAC3 protein (Ye, 2008), it exerts a positive effect on the transcriptional activity of NOTCH (Ando et al., 2003).

Altogether these observations support the hypothesis that the modulation of NFKB could potentially restore, in *mdx* FAPs, sensitivity to the NOTCH-dependent adipogenic signals. Given that NFKB is activated by inflammatory stimuli and FAPs from *mdx* mice are chronically exposed to inflammatory cytokines, we asked whether an inflammatory environment had any effect on *mdx* FAPs differentiation.

### 6. Synergic effect of muscle inflammatory response and NOTCH pathway in the regulation of *mdx* FAPs differentiation

Prompted by the observation that the *mdx* proteome is perturbed in key mediators of the crosstalk between the inflammatory response, the NOTCH and the adipogenic pathways, we asked whether an inflammatory environment had any effect on FAP differentiation. To address this question we isolated FAPs from young *mdx* mice and treated them with conditioned media obtained from CD45^+^ cells from muscles of young wt, *ctx* and *mdx* mice (Fig. 7 A). Such conditioned medium was then used to treat FAPs. We could not observe any significant difference in adipogenic differentiation in cells treated with CM from all three conditions (wt, *ctx, mdx*), compared to untreated controls. However, when *mdx* FAPs treated with CM were seeded on the NOTCH ligand, we could observe significant adipogenic inhibition in all three conditions. This suggests a cooperation of some factor produced by inflammatory cells and the NOTCH ligand in the negative regulation of *mdx* FAPs differentiation *ex vivo* (Fig. 7 B-E).

**Figure 7.**
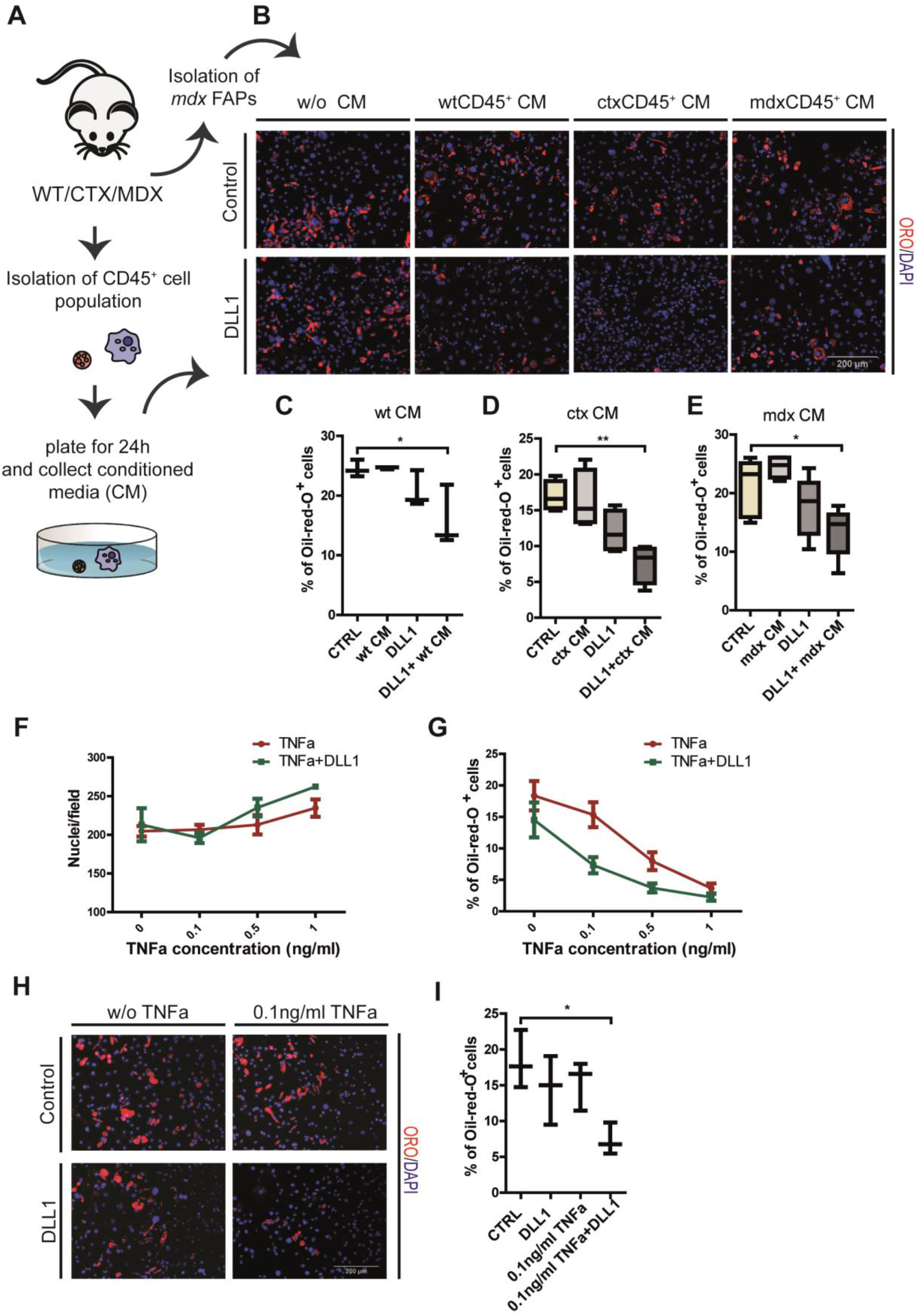
Signals secreted from inflammatory cells relieves the *mdx* FAP insensitivity to inhibition of adipogenesis mediated by NOTCH activation. A) Schema of the experimental procedure. (B) FAPs were isolated from *mdx* mice, seeded on plates containing IgG (control) or DLL1 and treated every 48h with conditioned medium obtained from CD45^+^ cells from *wt, ctx* and *mdx* mice. In order to obtain CD45^+^ conditioned media (CM), the CD45^+^ cell fraction was seeded for 24h in RPMI medium containing 10% FBS. After 24h the conditioned medium was collected, filtered to remove unattached cells and debris, and stored at 4 degrees. Adipocytes were stained with Oil red O and nuclei with DAPI. (C,D, E) The percentages of adipocytes are presented as box plots of at least three different experiments. FAPs±DLL1 were treated with 0.1, 0.5 and 1 ng/ml TNF-a and (F) average number of nuclei and (G) percentage of adipocytes were quantified. Data are represented as average ± SEM (n=3). (H) Representative immunofluorescence images of adipogenesis treated of FAPs treated with 0.1ng/ml TNF-a in the presence or not of the NOTCH ligand DLL1. Adipocytes were stained with Oil red O and nuclei with DAPI. (I) The percentage of adipocytes is presented as a box plot (n=3). Box plots show median and interquartile range with whiskers extended to minimum and maximum values. All quantifications were done using the CellProfiler software. Statistical significance was evaluated by the ANOVA test (*p<0.05, **p<0.01).

Tumor necrosis factor-alpha (TNF-a) is a pro-inflammatory cytokine expressed by macrophages type 1 during the inflammatory phase of the regeneration process (Tidball and Villalta, 2010). Its expression reaches a peak around day 3 post-injury (Lemos et al., 2015). We investigated whether this cytokine could be held responsible for the observed effect. FAPs were isolated from young *mdx* mice and treated with different concentrations of TNF-a every 48h for 8 days (Fig. 7 F-G). We observed a dose dependent inhibition of adipogenic differentiation. No apoptotic effect mediated by TNF-a was detected up to concentrations as high as 10 ng/ml (not shown). Interestingly even low concentrations of TNF-a (0.1 ng/ml), which *per se* have little effect on *mdx* FAP adipogenesis, when combined with activation of the NOTCH pathway restore sensitivity to the anti-adipogenic effect. (Fig. 7 H, I). These results parallel those obtained with CD45^+^conditioned media, suggesting that of TNF-a and NOTCH cooperate synergistically to inhibit *mdx* FAPs differentiation.

## Discussion

Fibro-adipogenic progenitors (FAPs) play an important role in muscle regeneration by promoting satellite cells activation and differentiation (Joe et al., 2010, Heredia et al., 2013, Mozzetta et al., 2013). However, in pathological conditions they cause fibrosis and ectopic fat deposition (Uezumi et al., 2010, Uezumi et al., 2011, Lemos et al., 2015). The recognition of the role played by this cell population both in physiological regeneration and in pathological degeneration has grown in recent years.

FAPs are characterized as mesenchymal progenitor cells expressing the Pdgfra and SCA1 antigens. These surface markers identify a heterogeneous cell population that readily differentiates into adipocytes even in the absence of adipogenic stimuli, when isolated from the muscle microenvironment (Joe et al., 2010, Uezumi et al., 2010). Why this differentiation potential is not manifest *in vivo* in the healthy muscle is still an unanswered question. These observations imply the existence of anti-adipogenic signals afforded by muscle resident cell types that, in physiological conditions, restrain the FAP adipogenic potential. A second important question relates to the reasons for the failure in the mechanisms that controls FAP adipogenesis in disease conditions, as observed for instance in DMD patients or in other pathologies causing accumulation of intramuscular adipose tissue. In our work we considered cell autonomous mechanisms that would make, with time, FAPs from a chronically inflamed muscle less sensitive to anti adipogenic signals. To this aim we analyzed the cellular and molecular differences underlying the adipogenic potentials of FAPs from wild type and *mdx* mice, by looking at the perturbation of their protein expression profile and by analyzing, by deep proteomics, the perturbation of molecular pathways involved in regulation of adipogenesis.

### Single cell mass cytometry reveals cell state heterogeneity in FAPs from different backgrounds

Despite sharing a rather homogeneous differentiation potential, FAPs are characterized by a substantial degree of heterogeneity in the expression of FAP-specific surface markers when analyzed by single cell mass cytometry. For instance we observed two populations with significant quantitative difference in the expression of the SCA1 and CD34 antigens (Joe et al., 2010). The population with low expression levels was detected in wild type FAPs, while the population with high levels of both antigens included the majority of FAPs from the preparation of regenerating muscles after cardiotoxin treatment (*ctx*). The FAP preparations from *mdx* mice muscles contained both populations in approximately equal amounts. Considering that the three samples derive from diverse muscle conditions influencing FAPs activation, the different expression levels of SCA1 and CD34 could reflect different cellular states. In agreement with this observation, it has been documented that different levels of CD34 expression correlate with proliferation and quiescence of hematopoietic stem cells *in vitro* (Dooley et al., 2004), while SCA1 expression has been linked to control of proliferation, self-renewal and differentiation of myoblasts and mesenchymal stem cells (Mitchell et al., 2005, Epting et al., 2008, Bonyadi et al., 2003). Our results parallel these observations. Namely, *wt* FAPs, isolated from an unperturbed and quiescent stem cell niche, have low levels of CD34 and SCA1, in accordance with their inactive state, while *mdx* and *ctx* FAPs, experiencing a regenerative environment that stimulates a proliferative state, express higher levels of these two antigens.

In addition we identified several subpopulations, which are differently populated in wt, *mdx* and *ctx* FAP preparations. In particular we singled out a subpopulation (A in Fig. 2) characterized by the expression of the CD146 antigen. Interestingly, type 1 pericytes, beside common pericyte markers such as NG2 and CD146, also express SCA1 and CD140α on their surface and *in vitro* show adipogenic potential (Birbrair et al., 2013). Overall, these observations highlight an overlap in the profile of gene expression of FAPs and pericytes and point to the heterogeneity and complexity of muscle mononuclear cell populations.

### Regulation of NOTCH adipogenesis by activation of the NOTCH pathway

Single cell mass cytometry and proteomics profiling highlighted the diversity of FAPs from different genetic or regenerating backgrounds reflecting distinct cellular states induced by different muscle environments. Still, *in vivo,* FAP adipogenic differentiation is restrained in all conditions suggesting the existence of signals that negatively control adipogenesis. Evolutionarily conserved pathways such as Hedgehog, Wnt and NOTCH have been implicated in the regulation of adipogenesis in mesenchimal stem cells (Rosen and MacDougald, 2006). We looked at the effect of activators and inhibitors of these three pathways on the modulation of FAP adipogenesis and we identified NOTCH as the pathway whose perturbation had the clearest impact.

NOTCH signaling plays an important role in skeletal muscle homeostasis and regeneration by regulating satellite cell states: quiescence, activation, proliferation and differentiation (Bjornson et al., 2012, Conboy and Rando, 2002, Lin et al., 2013, Mourikis et al., 2012, Yin et al., 2013, Luo et al., 2005). Here we have shown that the suppression of NOTCH signaling via inhibition of γ-secretase stimulates FAPs differentiation in a dose dependent manner, while activation by the NOTCH ligand DLL1 leads to a significant inhibition of adipogenesis. The involvement of NOTCH in controlling adipogenesis has been reported in different cell types, including the 3T3-L1 cell line (an adipogenic cell lineage) and human mesenchymal stem cells (Song et al., 2015, Ross et al., 2004, Osathanon et al., 2012). Our results highlight that NOTCH also controls adipogenesis of muscle FAPs and add to our understanding of skeletal muscle physiology.

Consistently, *ctx*-treated mice suffered mild adipocyte accumulation in the myofiber interstitial area when treated with DAPT starting at day 3 post-injury. The observed limited adipocyte infiltrations after inhibition of NOTCH suggest that NOTCH is involved in controlling ectopic fat infiltrations *in vivo* while at the same time implying that additional regulatory mechanisms are likely to play a role in controlling FAP differentiation.

### FAPs from mdx mice have limited sensitivity to NOTCH inhibition of adipogenesis

Interestingly, FAPs isolated from young *mdx* mice are unresponsive to NOTCH mediated inhibition of adipogenic differentiation by the DLL1 ligand when cultivated *ex vivo.* This finding highlights an important difference between *mdx* and wild type FAPs, difference that could explain fat deposition in the muscles of ageing *mdx* mice. We hypothesized that the observed insensitivity to NOTCH might be induced by the inflammatory environment in the degenerating *mdx* muscle. Thus we also tested FAPs isolated from a cardiotoxin damaged wild type muscle, another model of muscle regeneration. Somewhat contrary to our expectation, FAPs isolated from *ctx*-treated wild type mice are inhibited in their adipocytic differentiation when the NOTCH pathway is activated by the DLL1 ligand.

Thus, FAPs from *mdx* mice are insensitive to NOTCH inhibition *ex vivo.* However, *in vivo* the FAPs from young *mdx* mice do not cause adipocytic infiltrations suggesting that adipogenesis in the muscle is under the control of multiple redundant mechanisms and that some additional form of negative control is still functional even when NOTCH is failing (McDonald et al., 2015, DiMario et al., 1991, Grounds et al., 2008). The change in the muscle environment in ageing would expose the NOTCH defect and cause fat deposition. NOTCH insensitivity of *mdx* FAPs is not the consequence of a defect in the classical NOTCH pathway. In fact NOTCH signaling is active in freshly isolated *mdx* FAPs, as revealed by the expression of HES-1, product of a NOTCH target gene that is significantly higher in *mdx* FAPs, and comparable with that observed in FAPs from *ctx*-treated muscles.

### Deep proteomics reveals pathways that are perturbed in *mdx* FAPs

To further investigate the molecular mechanisms underlying the phenotypic differences observed in wt, *ctx* and *mdx* FAPs, we performed quantitative MS-based proteomic studies. Employing the recently developed ‘proteomic ruler’ method (Wisniewski et al., 2014)–we estimated protein copy numbers across the FAP proteomes. By this approach we obtained a global picture of the proteome-wide remodeling occurring in FAPs from wild type and regenerating muscles (young dystrophic and cardiotoxin-treated mice). Additionally, our approach enabled the identification of several hundred proteins significantly modulated in the *mdx* and *ctx* background. Here we focus on the analysis of the mechanisms that are perturbed in mdx FAPs and may affect control of adipogenesis. However, our dataset will serve as a more general resource for hypothesis-driven research to investigate the molecular mechanisms underlying functions that are perturbed in FAPs from dystrophic organisms.

Overlaying MS-based proteomic profiles onto literature-derived signaling pathways has been used to extract key mechanistic information (Sacco et al., 2016a, Sacco et al., 2016b). By this approach, we drew a network of adipogenesis-related pathways that are perturbed specifically in *mdx* FAPs and identified potential mechanisms which may explain the different sensitivity of *mdx* FAPs to anti-adipogenic NOTCH signals. Here we focused on the upregulation of proteins involved in retinol metabolism exclusively in *mdx* FAPs, and in particular of RXRA receptor, which is known to form a complex with PPARg to positively regulate adipogenesis (de Vera et al., 2017).

Our network analysis highlighted the central role of the NFKB signaling pathway that is activated in response to inflammatory stimuli such as TNF-a and is known to inhibit the activity of RXRA-PPARg while enhancing NOTCH signaling (Maniati et al., 2011, Ando et al., 2003, Ye, 2008).

### Synergic anti-adipogenic cross talk between inflammation and NOTCH signaling

Considering that *mdx* mice are chronically exposed to inflammation, we further investigated the involvement of inflammatory signals in modulating *mdx* FAPs differentiation. Interestingly, *mdx* FAPs treated with conditioned media from hematopoietic cells (CD45^+^ cell fraction) reacquire their sensitivity to NOTCH and their adypocite differentiation is significantly reduced. The hematopoietic CD45^+^ cells are purified from muscles 72 hours post-injury when TNF-a secretion from macrophages peaks (Lemos et al., 2015). We hypothesized that this cytokine could drive the observed synergic effect with NOTCH signaling. A cross talk between TNF-a and NOTCH signaling has been already reported in a variety of cell types (Jiao et al., 2012, Fazio and Ricciardiello, 2016, Ando et al., 2003, Maniati et al., 2011). We observed that physiological concentrations of TNF-a (Abdel-Salam et al., 2009), synergistically cooperate with NOTCH in the regulation of FAP adipogenesis. Overall, our results support a model whereby in young dystrophic muscles inflammatory stimuli, released by hematopoietic cells, boost the NOTCH signaling function and likely decrease RXRA-PPARg transcriptional activity, consequently restraining the adipogenic potential of FAPs. In our model, in young *mdx* mice, FAPs are insensitive to NOTCH anti-adipogenic signals, but do not cause adipocyte infiltrations *in vivo* thanks to the sustained inflammatory environment (Porter et al., 2002). We have observed a substantial decrease in macrophage infiltrations in muscle of old *mdx* mice (unpublished), supporting the idea that with ageing, the decrease in inflammation unveils the flaw in NOTCH signaling thus making adipogenesis control insufficient to prevent fat deposition.

Our results support a novel role of NOTCH signaling in skeletal muscle homeostasis. Here we propose a new molecular mechanism to explain fat infiltrations during disease progression. These findings may pave the way to the development of new therapeutic strategies aimed at ameliorating the disease phenotype in muscular dystrophies.

## Author contributions

Conceptualization, Mi.Ma. F.S. and G.C.; Investigation Mi.Ma., F.S.,L.L.P., F.S.,T.P. and C.F.; Data Curation, E.M. Writing-Original Draft, Mi.Ma. and G.C., Writing Review & Editing F.S., L.C. and C.G.; Funding Acquisition, G.C., Ma.Ma.; Supervision G.C., L.C. C.G. Ma.Ma.

## Acknowlegments

This work was supported by the DEPTH project of the European Research Council (grant agreement 322749) to GC.

## Materials and methods

### Contact for Reagent and Resource Sharing

Further information and requests for resources and reagents should be directed to and will be fulfilled by the Lead Contact, Gianni Cesareni.

### Experimental Model and Subject Details

#### Mice

In all the experiments young (6 weeks old) wild-type C57BL/6 mice or C57BL/6ScSn-Dmd^mdx^/J (*mdx* mice) purchased from the Jackson Laboratories were used. Mice were maintained according to the standard animal facility procedures and experiments on animals were conducted according to the rules of good animal experimentation I.A.C.U.C. n°432 of March 12 2006.

For cardiotoxin and glycerol muscle injury, wild-type C57BL/6 mice were first anesthetized with an intramuscular injection of physiologic saline (10 ml/kg) containing ketamine (5 mg/ml) and xylazine (1 mg/ml) after which 10 μM of cardiotoxin (Latoxan L81-02) or 20 μl of 50% v/v glycerol (glycerol + Hbss) were administered intramuscularly into the tibialis anterior (TA), quadriceps and gastrocnemius (GC). From day 3 post-injury (PI) mice were treated 3 times a week with intraperitoneal injection of the γ-secretase inhibitor, DAPT (95% corn oil/5% DAPT, 30 mg/kg, Sigma Aldrich) until the 21st day PI. Control mice were treated with vehicle (95% corn oil/5% DMSO). All experimental protocols were approved by the internal Animal Research Ethical Committee according to the Italian Ministry of Health regulation.

### Method Details

#### Cell isolation and MACS separation procedure

For preparation of mononuclear cell suspension, hind limbs were isolated from mice, minced mechanically and then subjected to an enzymatic dissociation for 1h at 37°C. The enzymatic mix contained 2μg/ml collagenase A, 2,4U/ml dispase II and 0,01mg/ml DNase I diluted in D-PBS with Calcium and Magnesium. Upon dissociation the enzymatic reaction was stopped with Hank’s Balanced Salt Solution with calcium and magnesium (HBSS Gibco, cat#14025-092) supplemented with 0.2% Bovine serum albumin (BSA) and 1% Penicillin-Streptomycin (P/S, 10.000 U/ml). The cells suspension was subjected to sequential filtration through a 100 μm, 70 μm and 40 μm cell strainer and centrifugations at 600 x g for 5 min. The lysis of red blood cells was performed with RBC Lysis Buffer (Santa Cruz Biotechnology).

Satellite cells and FAPs were isolated from the heterogeneous cell suspension using MACS microbeads technology. For magnetic beads separation the micro beads conjugated antibodies against CD45, CD31, α7-integrin and SCA1 were used. All antiboides were purchased from Miltenyi biotech. Satellite cells were selected as CD45^-^CD31^-^α7-integrin^+^ cells and FAPs as CD45^-^CD31^-^α7-integrin^-^SCA1^+^.

#### Cell culture conditions

FAPs were cultured at 37°C and 5% CO_2_ in growth medium containing Dulbecco modified Eagle medium (DMEM) supplemented with 20% heat-inactivated fetal bovine serum (FBS), 1% penicillin-streptomycin, 1mM sodium pyruvate and 10mM HEPES.

Satellite cells were cultured at 37°C and 5% CO2 on gelatin-coated plates in DMEM supplemented with 20% FBS, 10% heat-inactivated horse serum (HS), 2.5 ng/ml bFGF (PeproTech, cat#450-33), penicillin-streptomycin, 1mM sodium pyruvate and 10mM HEPES. For co-culture conditions cells were seeded in 1:1 ratio. For indirect co-cultures the cell culture inserts, transwells (Croning, cat#353104), with 1μm pore were used for 24-multiwell plates. Freshly isolated FAPs were plated on porous membrane of the transwell, while satellite cells were seeded on the bottom of the plate previously coated with gelatin.

In all experiments cell were seeded in 96-multiwell plates (3000 cells per well) or in 24-multiwell plates (18 000 cells per well).

For adipogenic induction two different media were used. Adipogenic induction medium (AIM) consisted of DMEM supplemented with 20% FBS, 0.5 mM 3-isobutyl-1-methylxanthine (IBMX) (Sigma, cat#I5879), 0.4 mM dexamethasone (Sigma, cat#D4902) and 1 μg/ml insulin (Sigma, cat#I9278). Cells were maintained for 48h in AIM, which was then replaced with the adipogenic maintenance medium (AM) consisting of DMEM supplemented with 20% FBS and 1 μg/ml insulin.

For fibrogenic induction cells were treated with 10ng/ml of TGFb (PeproTech, cat#100-21) For activation or inhibition of NOTCH, WNT and Hedghog pathways the following reagents were used: DAPT (Sigma Aldrich, cat#D5942), DLL1-Fc (RnD systems cat# 5026-DL), Wnt10b (RnD systems cat# 2110-WN-010), DKK1 (RnD systems cat# 5897-DK-010), Smoothened Agonist (SAG) (Selleck Chemicals, cat#S7779), Itraconazol (Sigma Aldrich, cat#I6657).

For CD45^+^ conditioned media (CM), the CD45^+^ cell fraction was seeded for 24h in Roswell Park Memorial Institute (RPMI) 1640 Medium containing 10% FBS and 1% penicillin-streptomycin. After 24h the conditioned medium was collected, filtered to remove unattached cells and debris, and stored at 4 degrees.

#### Inhibition and activation of NOTCH signaling

For the inhibition of NOTCH signaling the γ-secretase inhibitor, N-[N-(3, 5-difluorophenacetyl)-L-alanyl]-S-phenylglycine t-butylester (DAPT) was purchased from Sigma Aldrich and dissolved in DMSO. Depending on experimental conditions cells were treated with different concentrations (2uM, 5uM and 10 uM) of DAPT every 48h, for 8 days. The control samples were treated with an equal quantity of DMSO.

For activation of NOTCH signaling cells were cultured in 96-multiwell plates previously coated with 50 μl/well of 10 μg/ml DLL1-Fc or control IgG2A-Fc, dissolved in PBS for 2h at room temperature (RT).

#### Immunofluorescence staining

Cell cultures were fixed with 2% paraformaldehyde (PFA) for 15 minutes. Cells were permeabilized in 0.1% Triton X-100 for 5 min, blocked with PBS containing 10% Serum 0, 1% and TritonX-100 in for 1h at RT and incubated with the primary antibody for 1h at RT. Cells were then washed three times and incubated with the corresponding secondary antibody for 30min at RT. Nuclei were counterstained with 1 mg/ml of 4’, 6-diamidino-2-phenylindole (DAPI) for 5 minutes at RT.

Cryosections were permeabilized with 0.3% TritonX-100 in PBS for 30 min RT, blocked with PBS containing 0.1% Triton X-100, 10% goat serum and 1% Glycin for 2h at RT, and incubated with the primary antibody overnight at 4°C. Samples were then washed and incubated with secondary antibody for 1h at RT. Nuclei were counterstained with DAPI.

The following antibodies were used: mouse anti-MYHC (1:2), rabbit anti-perilipin (1:100), anti-PPARg (1:200), rabbit secondary antibody Alexa Fluor 488 conjugated (1:250) and antimouse secondary antibody Alexa Fluor 488 conjugated (1:250). Images were acquired automatically with a LEICA fluorescent microscope (DMI6000B).

#### Oil red O and Sudan black staining

The Oil red O solution (Sigma Aldrich) was used for detection of lipid droplets in adipocytes in cell cultures. The stock solution (0.5% filtered solution of Oil red O in isopropanol) was dissolved in ddH20 in 3:2 ratios and filtered. The cells were incubated for 5 min at RT, followed by two washings with PBS and DAPI staining.

For detection of intramuscular fat infiltrates the Sudan Black B Lipid Stain Kit (GeneCopoeia, cat# VB3102) was used according to protocol provided by manufacturer.

#### Immunoblotting

Whole cell proteins were extracted with RIPA lysis buffer (150 mm NaCl, 50 mm Tris-HCl, 1% Nonidet P-40, 0.25% sodium deoxycholate) supplemented with 1 mM pervanadate, 1 mM NaF, protease inhibitor cocktail 200X (Sigma), inhibitor phosphatase cocktail I and II 100X (Sigma). Protein concentrations were determined by Bradford colorimetric assay (Bio-Rad). Total protein extracts (30μg) were separated by SDS-PAGE, transferered to membranes and incubated in blocking solution (5% milk and 0.1% Tween-20 in PBS) for 1h RT. Membranes were then incubated with primary antibodies overnight at 4°C. The membranes were then washed three times and incubated with anti-mouse or anti-rabbit secondary antibody conjugated with HRP (1:2500) for 1 hour at room temperature. The blots were visualized with an enhanced chemiluminescent immunoblotting detection system. The antibodies used were as follows: anti-MYHC (1:1000), rabbit anti-PPARg (1:1000,), rabbit anti-HES1 (1:1000), mouse anti-vinculin (1:1000), rabbit anti-actin (1:1000). Densitometric analysis was performed using ImageJ software.

#### xMAP Technology

The xMAP Technology was used for detection of cytokines, growth factors and chemokines in secretome of satellite cells, FAPs and co-cultures. Luminex assay Mouse Premixed Multi-Analyte Kit was purchased from RnD systems. The preparation of samples, standards, microparticle cocktails (antibody mix) and reagents were performed according to manufacturer’s instructions. The samples were read in technical duplicate by Luminex instrument (MAGPIX 4.2). In this work only data obtained for adiponectin expression levels are presented.

#### Single cell mass cytometry

For mass cytometry analysis in order to eliminate magnetic beads FAPs were isolated by two step indirect labeling using PE conjugated antibody against SCA1 and Anti-PE MultiSort Kit, which allows removal of magnetic beads. Both antibody and kit were purchased from Miltenyi biotec.

Freshly isolated FAPs were suspended in D-PBS without calcium and magnesium and incubated with Cell-ID cisplatin (Fluidigm, Cat# 201064) at a final concentration of 5 μM for 5 minutes at RT. The staining was quenched by adding the Maxpar Cell Staining Buffer (Fluidigm, Cat# 201068). Cells stained with the cisplatin were then barcoded and labelled for the detection of cell surface and intracellular antigens according to manufacturer instructions. Briefly, for barcoding cells were incubated with appropriate barcodes resuspended in 800 μl of Barcode Perm Buffer and incubated for 30 min at RT. Upon the incubation cells were centrifuged and washed twice with Maxpar Cell Staining Buffer and was proceeded to staining protocol.

Cells suspended in Maxpar Cell Staining Buffer were incubated for 30 min at RT with the cell surface protein antibody cocktail (final dilution of 1:100 for each antibody). The antibody cocktail contained: anti-SCA1, anti-CD34, anti-CD146, anti-CD140b, anti-CXCR4, anti-CD31, anti-Vimentin. For intracellular antigen labelling cells were fixed with 1x Maxpar Fix I Buffer (5x, Fluidigm, Cat# 201065) for 10 minutes at RT and then permeabilized with 4°C ultra-pure methanol (Fisher Scientific Cat# BP1105-4) for 15 minutes on ice. Cell were then incubated with intracellular protein antibody cocktail for 30 min at RT. The antibody cocktail contained: anti-Caspase3, anti-pSTAT1, anti-pSTAT3, anti-pCreb, anti-TNF-a After cell surface and intracellular staining, cells were labeled Iridium DNA intercalator. Cells were suspended and incubated for 1 hour at RT in the intercalation solution, composed of Cell-ID Intercalator-Ir (Fluidigm, Cat# 201192A, 125 μM) diluted 1:1000 with Maxpar Fix and Perm Buffer (Fluidigm, Cat# 201067) to a final concentration of 125 nM.

Prior to analysis the cell concentration was adjusted with Mili-Q water to 2.5–5 × 10^5^ cells/ml. The cell suspension was then filtered with 30 μm-cell strainer cap into 5 ml round bottom polystyrene tubes. Data were acquired using mass cytometry platform, of DVS Sciences (CyTOF2).

#### Total proteome sample preparation and MS analyses

All samples were lysed in GdmCl buffer, boiled and sonicated, as previously described. Proteins were digested using LysC and trypsin (1:100), overnight at 37°C, in digestion buffer (20mM Tris-HCl, pH 8.5, and 10% acetonitrile). The obtained peptides were desalted on C18 StageTips

#### LC-MS/MS measurements

Peptides were loaded on a 50 cm reversed phase column (75 μm inner diameter, packed inhouse with ReproSil-Pur C18-AQ 1.9 μm resin [Dr. Maisch GmbH]). An EASY-nLC 1000 system (Thermo Fisher Scientific) was directly coupled online with a mass spectrometer (Q Exactive Plus, Thermo Fisher Scientific) via a nano-electrospray source, and peptides were separated with a binary buffer system of buffer A (0.1% formic acid [FA]) and buffer B (80% acetonitrile plus 0.1% FA), at a flow rate of 250nl/min. Peptides were eluted with a gradient of 5-30% buffer B over 240 min followed by 30-95% buffer B over 10 min, resulting in approximately 4 hr gradients. The mass spectrometer was programmed to acquire in a data-dependent mode (Top15) using a fixed ion injection time strategy. Full scans were acquired in the Orbitrap mass analyzer with resolution 60,000 at 200 m/z (3E6 ions were accumulated with a maximum injection time of 25 ms).

### Quantification and Statistical Analysis

#### Analysis of mass cytometry data

For Mass cytometry data analysis we exploited viSNE software, a computational approach suitable for the visualization of high-dimensional single-cell data, based on t-Distributed Stochastic Neighbor embedding (t-SNE) algorithm. viSNE is used to represent single cells in a two dimensional plot (i.e, viSNE map). In particular, each cell is represented by a point in the high-dimensional space and each cell coordinate is the expression level (intensity) of one protein measured by mass cytometer. Applying viSNE alghoritm, high-dimensional space is projected into two dimensional space preserving pairwise distances between points. The cells expressing similar levels of monitored antigens are located closely on a bi-dimensional map. In addition, a third dimension is visualized through the color, so the expression of a parameter is displayed into the viSNE map as a gradation of color of events reported (Amir el et al., 2013).

#### Proteome data processing and analysis

Raw MS data were processed using MaxQuant version 1.5.3.15 (Cox and Mann, 2008; Cox et al., 2011) with an FDR < 0.01 at the level of proteins, peptides and modifications. Searches were performed against the Mouse UniProtKb FASTA database (2015). Enzyme specificity was set to trypsin, and the search included cysteine carbamidomethylation as a fixed modification and N-acetylation of protein, oxidation of methionine, as variable modifications. Up to three missed cleavages were allowed for protease digestion, and peptides had to be fully tryptic. Quantification was performed by MaxQuant, ‘match between runs’ was enabled, with a matching time window of 1 min. Bioinformatic analyses were performed with Perseus (www.perseus-framework.org). Significance was assessed using two-sample student’s t-test, for which replicates were grouped, and statistical tests performed with permutation-based FDR correction for multiple hypothesis testing. Proteomics raw data have been deposited in the ProteomeXchange Consortium (http://proteomecentral.proteomexchange.org) via the PRIDE partner repository with the data set identifier.

#### xMAP data analysis

The secretome analysis with Luminex (Fig. 4D) was performed in biological duplicate (n=2), where n represents cells isolated from multiple mice in two distinct experiment. The samples were read in technical duplicate by Luminex instrument (MAGPIX 4.2) and analyzed with xPONENT 3.1 software.

#### Image analysis

For adipogenic differentiation quantification we exploited CellProfiler software (Carpenter et al., 2006), using pipelines established in our group. Average number of nuclei was determined automatically with Cell Profiler and adipocytes differentiation was measured as the ratio of nuclei surrounded by Oil red O staining or positive for PPARg and the total number of nuclei per field. Quantifications were obtained by averaging the signal from 25 images. Satellite cells differentiation into myotubes was quantified using CellProfiler software as ratio of MYHC covered area over the total area of the analyzed field.

Mass cytometry analysis (Fig. 1 and 2) was performed in biological triplicates (n=3) where n represents cells isolated from multiple mice. In order to obtain enough cells to perform the experiment, cells isolated from different mice were combined as one biological replicate: 15 *wt* mice divided in 3 biological replicates (5 mice per replicate) were used for *wt* FAPs, 6 *mdx* mice divided in 3 biological replicates (2 mice per replicate) were used for *mdx* FAPs and 3 mice treated with *ctx* were used for *ctx* FAPs. In proteomics analysis (Fig. 6) each biological replicate represents cells isolated from one mouse, n=3. In *in vivo* experiment (Fig.4 K) n represents number of mice used (n control =2, n DAPT= 2, n glycerol=1). In all *in vitro* experiments n represents cells isolated from one mouse, in at least three independent isolation procedures. Each condition was analysed in technical duplicate.

For statistical analysis Student’s t-test or ANOVA were calculated using GraphPad Prism software to determine significant differences between means in all experiments. Box plots show median and interquartile range with whiskers extended to minimum and maximum values. Bar graphs show mean values ± SEM. The differences were considered significant at *p<0.05, ** p<0.01, *** p<0.001.

